# Reduced binding associated with resistance to Vip3Aa in the corn earworm (*Helicoverpa zea*)

**DOI:** 10.1101/2023.07.07.548161

**Authors:** Dawson D. Kerns, Fei Yang, David L. Kerns, Scott D. Stewart, Juan Luis Jurat-Fuentes

## Abstract

Transgenic corn and cotton expressing Cry and Vip insecticidal proteins from the bacterium, *Bacillus thuringiensis* (Bt), have been a valuable tool for the management of lepidopteran pests. In 2019, a Vip3Aa-resistant strain of *Helicoverpa zea* (CEW-Vip-RR) was isolated from F_2_ screens of field populations in Texas. Characterizing the resistance mechanism in this strain is important for predicting the sustained efficacy of current commercial Bt traits and guiding the development of future transgenic traits. Resistance to insecticidal proteins in Bt traits is commonly associated with reduced toxin binding, with the exception of Vip3Aa resistance being associated to altered proteolytic processing in the insect host gut. Therefore, Vip3Aa protoxin processing was tested by incubation with midgut fluids from CEW-Vip-RR relative to a susceptible strain (CEW-SS). Finding no significant processing differences, alterations in Vip3Aa binding were tested by comparing binding of radiolabeled and biotinylated Vip3Aa toxin to midgut brush border membrane vesicles (BBMV) from CEW-Vip-RR and CEW-SS larvae. Specific Vip3Aa binding to CEW-Vip-RR BBMV in these experiments was consistently reduced when compared with CEW-SS BBMV. These results support that an altered Vip3Aa- receptor is associated with resistance in CEW-Vip-RR. Understanding this resistance mechanism could have important implications for resistance management decisions considering widespread Cry1 and Cry2 resistance in *H. zea* populations.

**IMPORTANCE:** *Helicoverpa zea* is a major crop pest in the United States that is managed with transgenic corn and cotton producing insecticidal proteins from the bacterium, *Bacillus thuringiensis* (Bt). However, *H. zea* has evolved widespread resistance to the Cry proteins produced in Bt corn and cotton, leaving Vip3Aa as the only plant incorporated protectant in Bt crops consistently providing excellent control of *H. zea*. The benefits provided by Bt crops will be substantially reduced if widespread Vip3Aa resistance develops in *H. zea* field populations. Therefore, it is important to identify resistance alleles and mechanisms that contribute to Vip3Aa resistance to ensure that informed resistance management strategies are implemented. This study is the first report of reduced binding of Vip3Aa to midgut receptors associated with resistance.

## INTRODUCTION

The corn earworm or bollworm, *Helicoverpa zea* Bodie (Lepidoptera: Noctuidae), is one of the most damaging crop pests in the United States, causing annual losses of $90 million dollars in cotton alone (1). Transgenic corn and cotton that produce insecticidal proteins from the bacterium, *Bacillus thuringiensis* (Bt), have been used in the U.S. to manage *H. zea* for over 20 years (2). These transgenic crops currently account for 84 and 89 percent of U.S. corn and cotton acreage, respectively (3). Adoption of Bt crops provides effective pest management as well as economic and environmental benefits, including reduced insecticide applications and decreased mycotoxin levels in corn (4–8). However, the high adoption rate of Bt crops increases selection pressure for resistance, currently the main threat to the sustained benefits of Bt crops.

To date, populations of *H. zea* in the U.S. have evolved practical resistance to the Cry1Ab, Cry1Ac, Cry1A.105 and Cry2Ab2 Bt proteins produced in transgenic corn and cotton (9–13). Consequently, Vip3Aa remains the only highly effective insecticidal Bt protein for management of *H. zea* (13–15). However, alleles for resistance to Vip3Aa have already been isolated from field populations (16, 17). Estimations of resistance allele frequency and trends of reduced susceptibility to Vip3Aa in *H. zea* populations suggest the development of widespread Vip3Aa resistance could be imminent (14, 16, 18).

In contrast to the well-studied mode of action of Cry proteins (19), aspects of the Vip3Aa mode of action remain unknown (20). The Vip3Aa protein is produced as a protoxin with a molecular mass of ∼89 kDa (20), which forms a pyramid-shape tetramer in solution (21). Upon ingestion, the protoxin is processed by gut proteases to a 62-66 kDa core toxin and a smaller peptide fragment of ∼22 kDa that remains linked to the toxin core (22, 23). Activation induces drastic remodeling that releases the N-terminus segment of the protein and leads to the formation of a four-helix coiled coil sufficiently long to reach and permeate the lipid bilayer (21). In *H. zea* larvae, activated Vip3Aa specifically recognizes binding sites on the midgut brush border membrane that are not shared with Cry1Ac or Cry2Ab2 proteins (24). Evidence suggests binding is conducive to Vip3Aa insertion and pore formation on the brush border membrane of susceptible insects (25), yet there is also evidence that Vip3Aa could be endocytosed and induce apoptosis (26–30). After enterocyte death, subsequent midgut epithelium degeneration facilitates bacterial movement to the main body cavity and eventual death of the insect from septicemia (31–33). While alterations in any of these steps could potentially result in resistance to Vip3Aa, the limited experimental evidence available associates high-level (400-2,000-fold) Vip3Aa resistance with altered proteolytic processing (34) and reduced levels of alkaline phosphatase (35). In contrast, most cases of high-level Cry resistance have been attributed to alterations in midgut receptors and toxin binding, with lower resistance levels often associated with alterations in protoxin processing (36).

In 2019, a laboratory strain of *H. zea* displaying >588-fold resistance to Vip3Aa (CEW- Vip-RR) was isolated from an F_2_ screen of field populations in Texas (18). This strain did not exhibit cross-resistance to Cry1Ac or Cry2Ab2 (37). The goal of this study was to provide a mechanistic description of resistance to Vip3Aa in the CEW-Vip-RR strain, as there is currently no data available on Vip3Aa resistance mechanisms in *H. zea*. Proteolytic processing and binding of Vip3Aa in CEW-Vip-RR were assessed and compared to a near-isogenic, susceptible (CEW-SS) *H. zea* strain. Results from this study increase our understanding of Vip3Aa resistance mechanisms and contribute to the identification of Vip3Aa resistance genes.

## RESULTS

### Larval gut pH measurements

No differences in pH between untreated CEW-SS and CEW-Vip-RR larvae were detected in the foregut, anterior midgut, central midgut, posterior midgut, or hindgut (df = 256; *P ≥* 0.990 for all). The anterior, central, and posterior midgut regions of untreated larvae did not significantly differ, with pH values (mean ± standard error) of 9.47 ± 0.15, 9.92 ± 0.11, and 9.58 ± 0.16, respectively. In comparison, the foregut and hindgut regions had significantly lower (*P* < 0.0001) pH values of 7.27 ± 0.14 and 7.48 ± 0.08, respectively.

After treatment with a lethal dose of Vip3Aa protoxin, Vip3Aa toxin, or Cry1Ac protoxin, the pH of all three midgut regions in CEW-SS larvae were significantly reduced (*P* < 0.0001) compared to untreated larvae (Figure 1A). The pH of the foregut was not altered after treatment, whereas the pH in the hindgut was only reduced after treatment with Cry1Ac (t = - 4.12; df = 256; *P* = 0.0269). In contrast, the midgut pH of CEW-Vip-RR larvae was only significantly reduced (*P* < 0.0001) after treatment with Cry1Ac protoxin, and neither Vip3Aa protoxin nor Vip3Aa toxin significantly affected pH in any midgut region (Figure 1B, *P* = 1.0000). Similarly, only Cry1Ac reduced the pH of the foregut (t = -4.40; df = 256; *P* = 0.0092), and Vip3Aa treatments did not have a significant effect on pH (*P* = 1.0000). None of the treatments significantly altered the hindgut pH in CEW-Vip-RR larvae (*P* > 0.7000).

**Figure 1.**
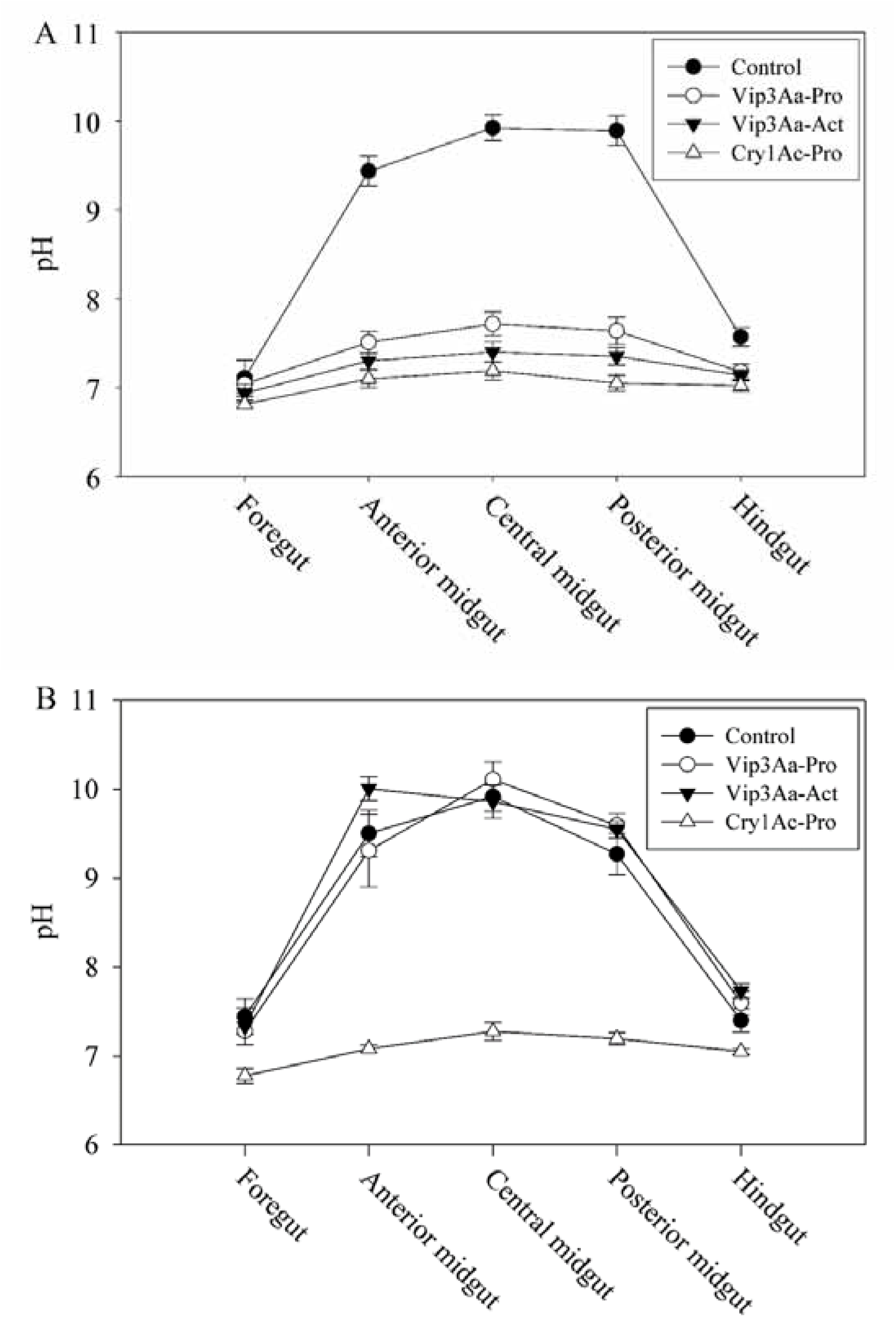
Gut pH measurements from five gut sections of CEW-SS (A) and CEW-Vip-RR (B) fourth-instar *H. zea* larvae. Groups of nine larvae were fed a droplet of sucrose water containing Vip3Aa protoxin (Vip3Aa-Pro), Vip3Aa activated toxin (Vip3Aa-Act), Cry1Ac protoxin (Cry1Ac-Pro), or with no treatment as a control (Control). Data plotted are the mean pH values and standard errors for each treatment in different gut regions.

### Vip3Aa and Cry1Ac protoxin processing

When incubated with *H. zea* larval gut fluids, Vip3Aa protoxin consistently took longer to process than Cry1Ac protoxin. Processing resulted in activated toxin forms of ∼65 kDa for Cry1Ac and ∼66 kDa for Vip3Aa (Fig. 2A). A band slightly below the band corresponding to the activated Vip3Aa toxin fragment was detected after the ∼89 kDa protoxin band had almost disappeared and became more predominant at the longest incubation period tested (21 hours). There were no differences in protein band profiles when comparing Vip3Aa or Cry1Ac protoxin processing with gut fluids from CEW-SS and CEW-Vip-RR larvae (Fig. 2A-B). There were also no differences in the rate of Vip3Aa and Cry1Ac protoxin processing by CEW-SS or CEW-Vip- RR gut fluids (t = 0.95; df = 4; *P* = 0.3940, Vip3Aa; t = 0.56; df = 4; *P* = 0.6044, Cry1Ac) (Fig. 2C-D).

**Figure 2.**
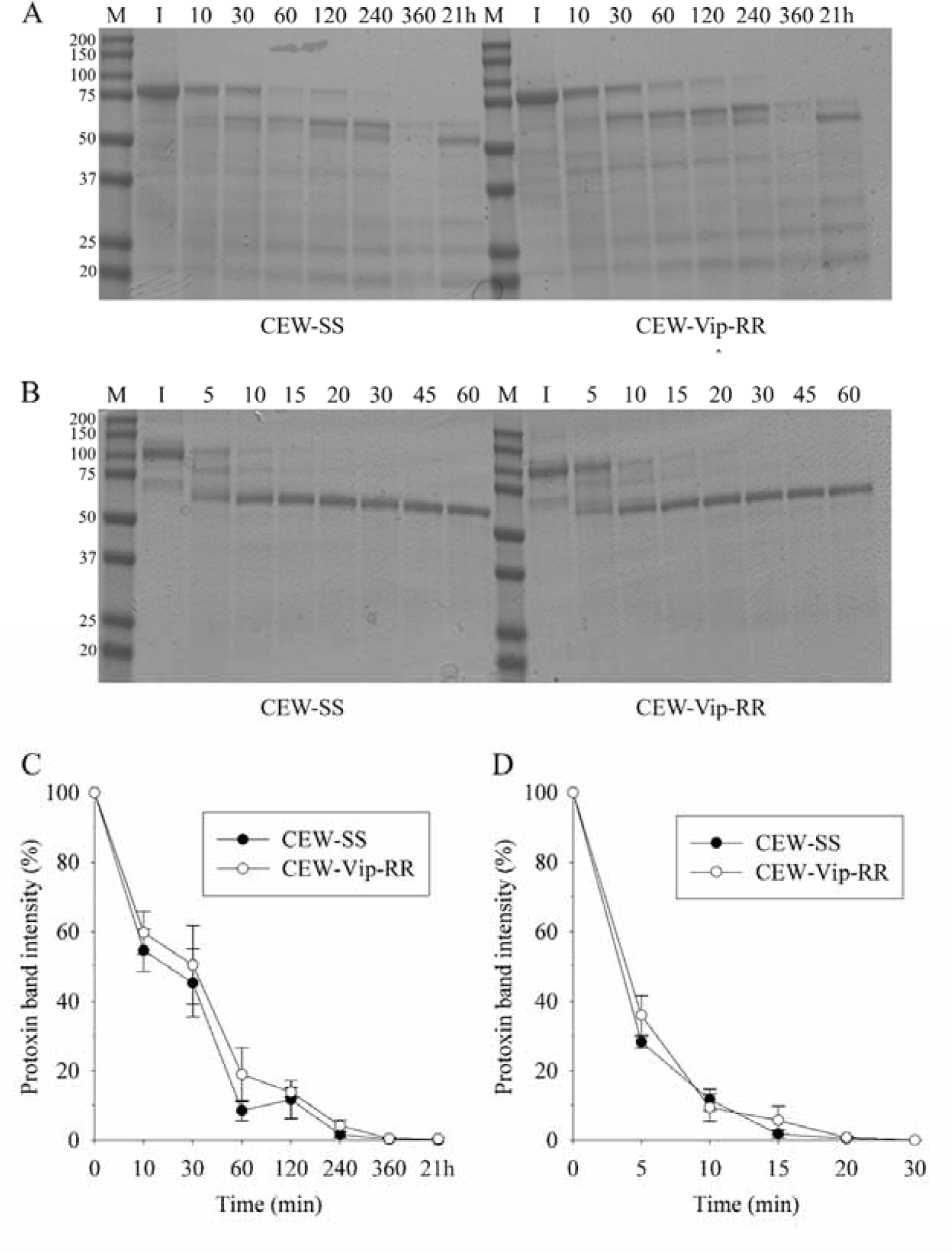
Incubation of Vip3Aa or Cry1Ac protoxin with midgut fluids extracted from CEW-SS and CEW-Vip-RR *H. zea* larvae. Protoxin was incubated with midgut fluid extracts and aliquots were taken at various timepoints. Vip3Aa (A) and Cry1Ac (B) samples were resolved using SDS-PAGE and stained with ProtoBlue safe; M: protein marker, I: Protoxin input. Percent Vip3Aa (C) and Cry1Ac (D) protoxin processing over time by midgut fluid extracts from CEW- SS and CEW-Vip-RR larvae. Plotted is the percent intensity of the protoxin bands for each timepoint relative to the intensity of the input protoxin band as estimated by densitometry. Each point and standard error bar are representative of three independent experiments. No differences in the rate of processing were detected (generalized linear mixed model; Tukey post-hoc; α= 0.05.

### Homologous competition assays with biotinylated Vip3Aa toxin

Western blots of binding assays with biotinylated Vip3Aa detected reduced total binding in BBMV from larvae of the CEW-Vip-RR strain compared to CEW-SS strain while levels of non-specific binding were similar in BBMV from both strains (Fig. 3B). Interestingly, the biotinylated Vip3Aa band detected in all binding reactions appeared slightly lower in size than the input Vip3Aa. The size of this Vip3Aa band did not correspond to any of the protein bands that appeared after 21 hours of processing Vip3Aa with gut fluids. Densitometry measurements detected similar intensity between the Vip3Aa band detected for total binding to CEW-SS BBMV and 10 ng of biotinylated Vip3Aa (Fig. 3B), suggesting that approximately three percent of the input biotinylated Vip3Aa toxin bound to the BBMV. On the other hand, approximately 2 ng of biotinylated Vip3Aa were estimated as bound to CEW-Vip3-RR larvae in total binding assays. Estimation of specific binding by subtracting non-specific from total binding determined a greater than six-fold reduction in specific binding of biotinylated Vip3Aa to CEW-Vip-RR compared to CEW-SS BBMV (t = 21.6; df = 6; *P* < 0.0001) (Fig. 3A).

**Figure 3.**
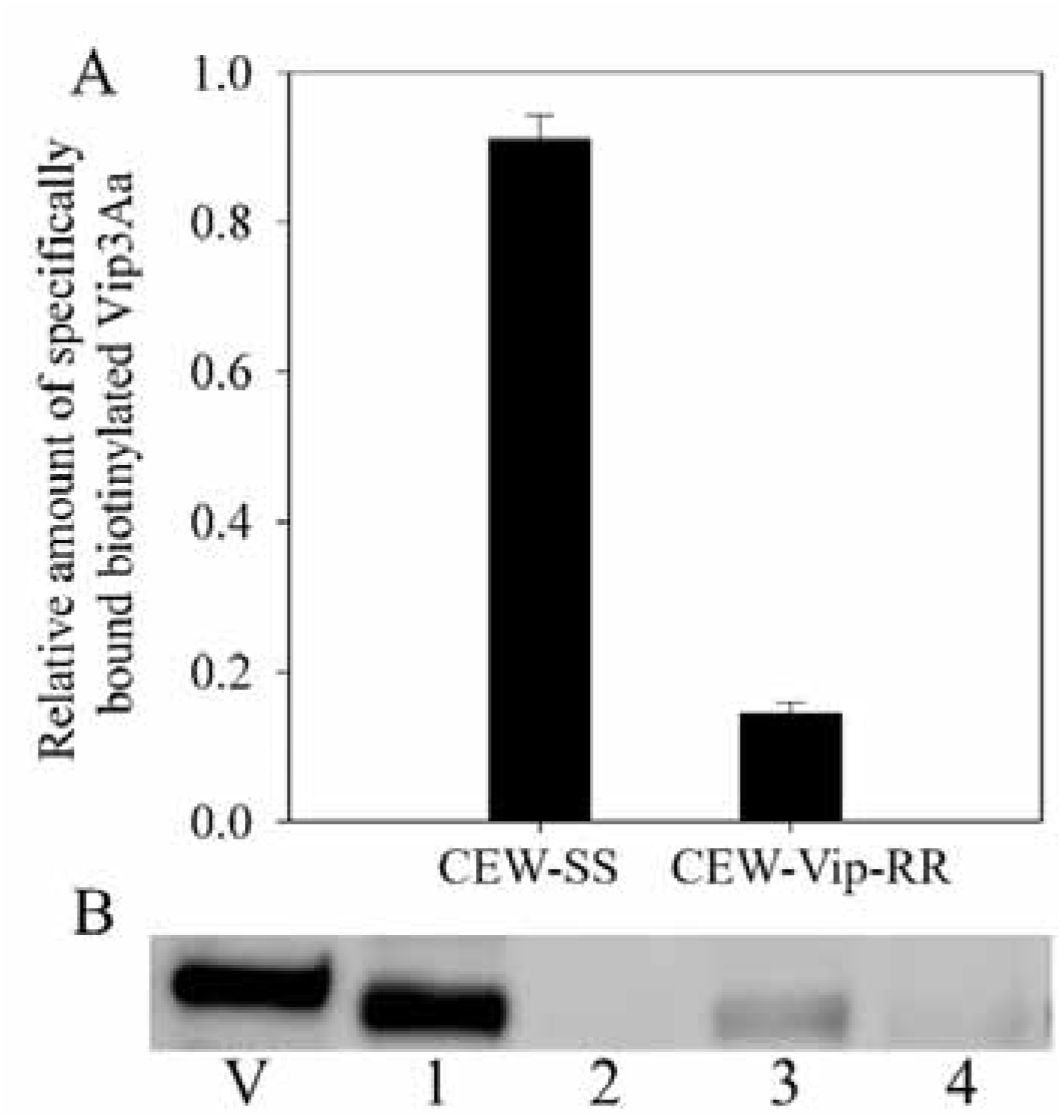
Binding of biotinylated Vip3Aa toxin bound to brush border membrane vesicles (BBMV) from CEW-SS and CEW-Vip-RR *H. zea* larvae. Brush border membrane vesicles were incubated with biotinylated Vip3Aa toxin with (non-specific binding) or without (total binding) a 100-fold molar excess of unlabeled Vip3Aa toxin. Bound biotinylated toxin was recovered by centrifugation and final pellets were suspended in 1X sample buffer before visualizing by western blotting (A); V: 10 ng biotinylated Vip3Aa toxin; 1: CEW-SS total binding; 2: CEW-SS non-specific binding; 3: CEW-Vip-RR total binding; 4: CEW-Vip-RR non-specific binding. Densitometry was used to estimate band intensity and specific binding was calculated by standardizing band intensity relative to the intensity of 10 ng of Vip3Aa toxin, then subtracting non-specific binding from total binding (B). Bars and standard errors are representative of the mean of two biologically independent experiments (different BBMV preparations) each with two technical replications. Specific binding of Vip3Aa was significantly reduced in CEW-Vip-RR relative to CEW-SS (independent t-test; α= 0.05).

### ^125^I-Vip3Aa and ^125^I-Cry1Ac binding assays

Saturation assays showed that BBMV from both strains bound Vip3Aa specifically, yet specific ^125^I-Vip3Aa binding was reduced in BBMV from CEW-Vip-RR compared to CEW-SS (t= -5.75; df = 41; *P* < 0.0001) (Fig. 4). Because specific Vip3Aa binding did not reach saturation, competitive binding assays were performed to better estimate binding parameters.

**Figure 4.**
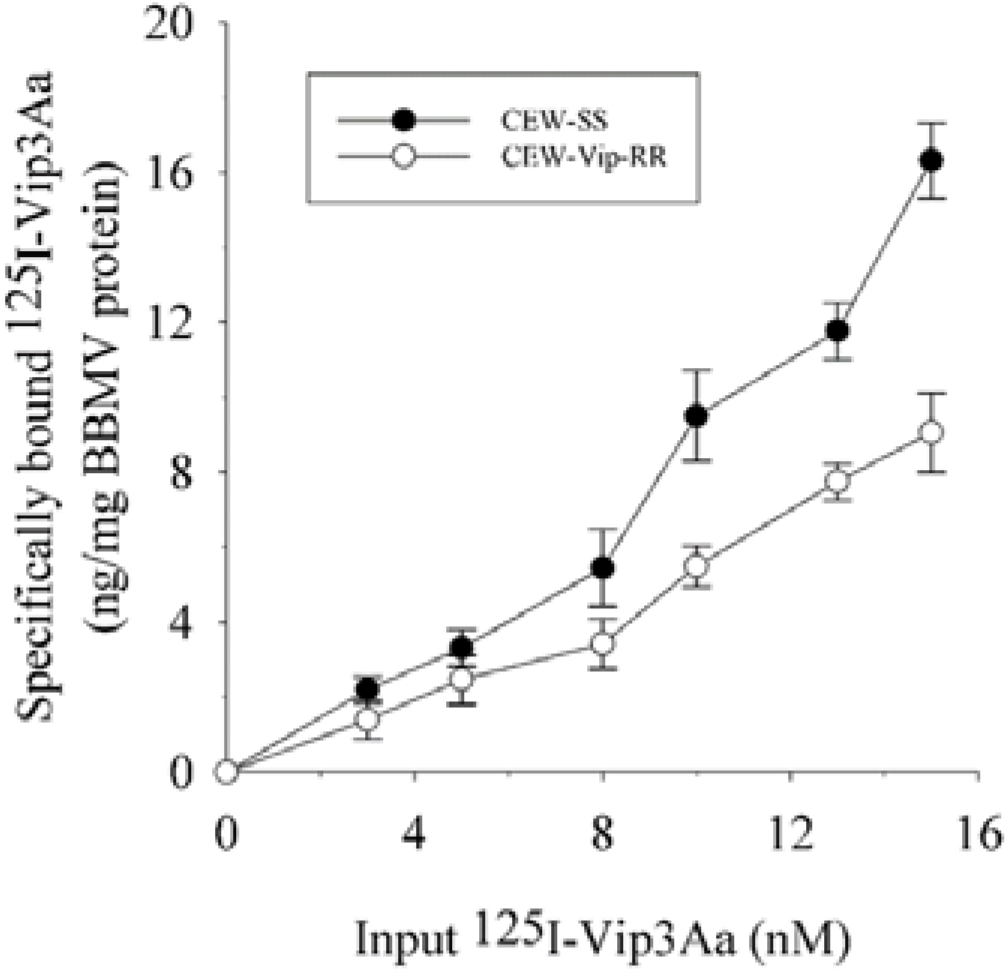
Specific binding of radiolabeled Vip3Aa toxin (^125^I-Vip3Aa) to brush border membrane vesicles (BBMV) from CEW-SS and CEW-Vip-RR *H. zea* larvae in saturation assays. Brush border membrane vesicles were incubated with increasing amounts of ^125^I-Vip3Aa with (non-specific binding) or without (total binding) a 100-fold molar excess of unlabeled Vip3Aa. Bound ^125^I-Vip3Aa was recovered by centrifugation and detected using a gamma counter. Specific binding was calculated by subtracting the amount of non-specifically bound ^125^I-Vip3Aa from the total bound ^125^I-Vip3Aa. Each point and standard error bar is representative of at least two biologically independent experiments (different BBMV preparations) each with two technical replications. Binding of ^125^I-Vip3Aa was significantly reduced in CEW-Vip-RR (generalized linear model; Tukey-Kramer post-hoc; α= 0.05).

Binding of ^125^I-Vip3Aa was displaced more effectively in BBMV from CEW-SS compared to CEW-Vip-RR at all tested unlabeled Vip3Aa concentrations (Fig. 5A). Thus, displacement reached 68% for CEW-SS, while the highest displacement for CEW-Vip3-RR was 51%. In contrast, no difference in competition of ^125^I-Cry1Ac binding was observed between BBMV from CEW-SS and CEW-Vip3-RR (Fig. 5B). Estimated binding parameters showed that Vip3Aa bound with >3.7-fold higher affinity to BBMV from CEW-SS than CEW-Vip-RR, while Cry1Ac binding parameters for BBMV from the two strains were similar (Table 1).

**Figure 5.**
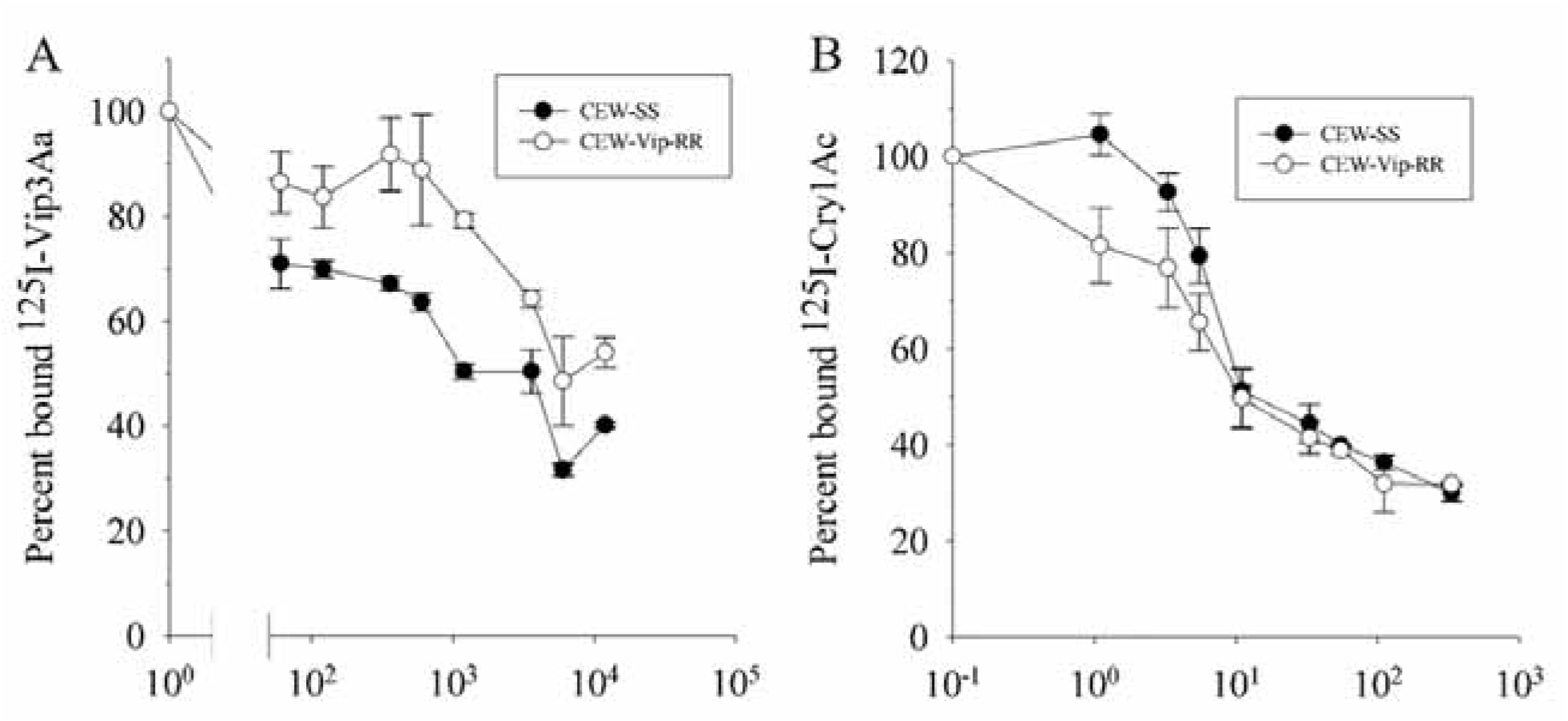
Competition assays of ^125^I-Vip3Aa (A) or ^125^I-Cry1Ac (B) ligands binding to brush border membrane vesicles (BBMV) from CEW-SS and CEW-Vip-RR *H. zea* larvae. Brush border membrane vesicles were incubated with radiolabeled toxin and increasing amounts of unlabeled homologous competitor. Amounts of competitor tested were selected on the basis of labeled toxin input and consistency of competition. Bound toxin was recovered through centrifugation, and radiolabeled toxin was detected by a gamma counter. Each point and standard error bar is representative of at least two biologically independent experiments (different BBMV preparations), each technically replicated two times.

**Table 1.** Equilibrium disassociation constants and binding site concentrations for Vip3Aa and Cry1Ac toxins from competition binding assays with BBMV from *H. zea* larvae.

| Protein | Strain<sup>a</sup> | Mean ± SEM | |
| --- | --- | --- | --- |
|  |  | K<sub>d</sub> (nM) | R<sub>t</sub> (pmol/mg) |
| Vip3Aa | CEW-SS | 61 ± 29 | 0.22 ± 0.03 |
| | CEW-Vip-RR | 228 ± 54 | 0.22 ± 0.02 |
| Cry1Ac | CEW-SS | 1.5 ± 0.57 | 0.3 ± 0.07 |
| | CEW-Vip-RR | 0.79 ± 0.24 | 0.3 ± 0.05 |

<sup>a</sup> CEW-SS, susceptible; CEW-Vip-RR, Vip3Aa resistant

### ALP and APN enzymatic activity assays

Given that ALP levels were reduced in Vip3Aa-resistant *H. virescens* (35), we measured enzymatic ALP and APN activity in midgut homogenates and BBMV. No differences in specific ALP enzymatic activity were detected between BBMV (t = -1.02; df = 10; *P* = 0.3298) or midgut homogenates (t = 0.716; df = 10; *P* = 0.4900) from larvae of the CEW-SS and CEW-Vip3-RR strains (Fig. 6). Similarly, there were no differences in specific APN activity in BBMV (t = 0.419; df = 10; *P* = 0.6841) or midgut homogenate (t = -0.516; df = 10; *P* = 0.6172) between BBMV from both strains.

**Figure 6.**
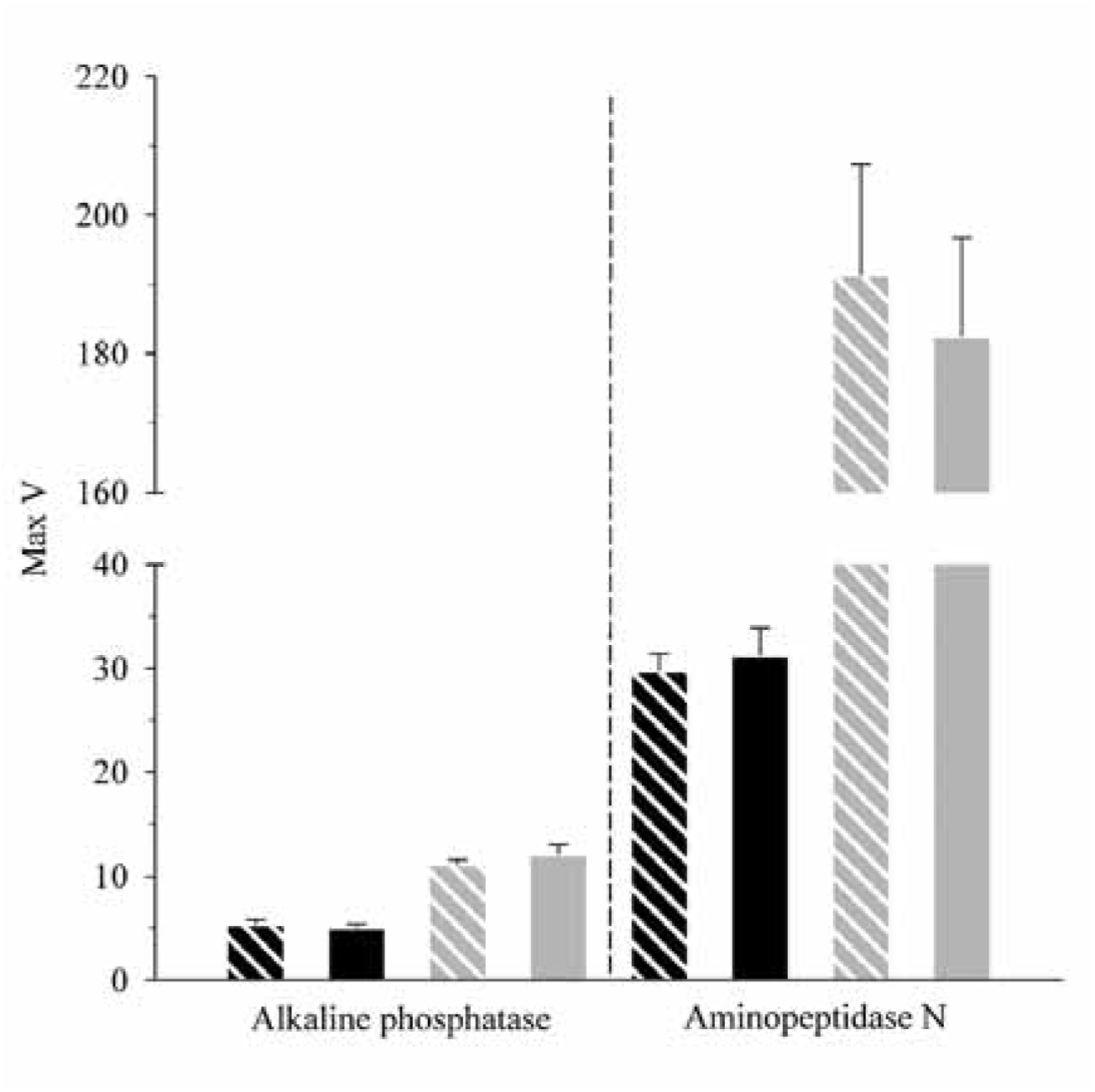
Alkaline phosphatase and aminopeptidase N activity from midgut homogenates and BBMV of two *H. zea* strains (black stripes: homogenate, CEW-SS; black solid: homogenate, CEW-Vip-RR; gray stripes: BBMV, CEW-SS; gray solid: BBMV, CEW-Vip-RR). Enzyme activity was measured in samples using leucine *p*-nitroanilide and *p*-nitrophenyl phosphate as substrates. Bars and standard errors represent three technical replications of two different BBMV preparations or their corresponding midgut homogenates. No significant differences in enzymatic activity were detected between CEW-SS and CEW-Vip-RR homogenates (independent t-test; α= 0.05).

## DISCUSSION

The extensive use of Bt corn and cotton in the U.S. has resulted in widespread practical resistance to Cry1A and Cry2A proteins in *H. zea* populations (9–13). Consequently, Vip3Aa is the sole plant incorporated protectant in Bt crops that consistently provides excellent control of *H. zea*. However, the identification of alleles associated with high levels of resistance to Vip3Aa in field *H. zea* populations raises concerns for practical resistance emergence if suitable resistance management strategies are not implemented promptly (15).

Understanding the molecular mechanisms underlying Vip3Aa resistance is critical to developing effective resistance management strategies. Data in this study support that high levels of resistance to Vip3Aa in a field-derived strain of *H. zea* are associated with reduced Vip3Aa binding to midgut receptors. This observation represents the first evidence supporting alterations in toxin binding as a mechanism of resistance to Vip3Aa. Additionally, the study identified that alterations in midgut pH, Vip3Aa processing, and ALP levels are not associated with Vip3Aa resistance in the CEW-Vip-RR strain.

Previous studies presented evidence that Vip3Aa binding and midgut proteolytic activity is highly dependent on pH (23, 38, 39). In contrast, no differences in midgut pH were detected between untreated larvae from the CEW-SS and CEW-Vip3-RR strains. Reduced midgut pH was observed after treatment with Cry1Ac or Vip3Aa protoxins and toxins in CEW-SS larvae. This phenomenon was previously observed in *H. zea* treated with Cry1Ac (40) and is likely a result of midgut ion homeostasis disruption after toxin damage to the gut epithelium (41). These observations suggest that Vip3Aa injury to the gut epithelium also has an effect on midgut homeostasis.

Lack of decreased midgut pH in CEW-Vip-RR larvae fed with either Vip3Aa protoxin or toxin supports that protoxin processing is not involved in resistance. This conclusion is further supported by the results from Vip3Aa and Cry1Ac processing experiments. Considering that alterations to proteolytic processing have often been associated with cross-resistance among Bt proteins that do not share binding sites (42), these observations align with the lack of cross-resistance to Cry1Ac in CEW-Vip-RR (38). However, these findings contrast with previous reports of slower processing associated with resistance to Vip3Aa in *Helicoverpa armigera* (34), and differences in processing kinetics being associated with distinct Vip3Aa toxicity (43, 44).

Despite these observations, evidence for reduced proteolytic processing as a major Vip3Aa resistance mechanism is limited because the observed reductions are either less than the amount expected to generate high resistance levels (34), or they do not result in high levels of Vip3Aa resistance (43, 44).

Alterations in binding are the most common resistance mechanism attributed to high levels of resistance to Cry1 proteins (36). However, previous reports did not detect alterations of Vip3Aa binding in Vip3Aa-resistant insects (34, 35, 45). Binding of ^125^I-Vip3Aa to BBMV from a laboratory-selected strain of *Chloridea virescens* with polygenic resistance to Vip3Aa did not differ from an unselected strain from the same initial population. Similarly, a laboratory-selected Vip3Aa resistant strain of *Mythimna separata* and a Vip3Aa-resistant *H. armigera* strain isolated from an F_2_ screen showed no alterations in ^125^I-Vip3Aa binding to BBMV, compared to reference susceptible strains. Results from binding assays in this study using biotinylated or radiolabeled Vip3Aa toxin support that binding to midgut brush border membrane receptors is reduced in CEW-Vip-RR relative to CEW-SS. Interestingly, this reduction was not absolute, and in contrast to some cases of resistance to Cry proteins (46–49). In fact, there are also cases of binding site alteration associated with resistant insect strains that resulted in only a partial reduction of binding when compared to a susceptible strain (50–52), including cases of practical resistance to Bt corn (53) and knockout of a Cry receptor (54). Considering that there is evidence that Vip3Aa binds to at least two sites in *Spodoptera spp.* (55, 56), it is plausible that the reduced binding competition detected represents lack of binding to a subpopulation of Vip3Aa binding sites in CEW-Vip-RR. As expected, given the lack of cross-resistance to Cry1Ac, no alterations in Cry1Ac binding were detected in BBMV from CEW-Vip-RR compared to CEW-SS.

The binding parameters obtained using radiolabeled Vip3Aa agreed with the binding affinity of this protein observed in other studies (23, 34, 35, 45). This lower affinity compared to Cry proteins may explain the lack of radiolabeled Vip3Aa binding saturation. Other mechanistic studies of Vip3Aa resistance reported higher concentrations of binding sites that did not correspond with reduced Vip3Aa binding (34, 35, 45). The detection of reduced Vip3Aa binding in this study in contrast to previous studies could be due to differences in experimental methodology and insect species tested. Binding of Vip3Aa is heavily influenced by conditions in the binding reaction (23), thus the binding conditions in this study may have been more favorable for detecting binding site alterations. Additionally, the differences in binding between CEW-SS and CEW-Vip-RR BBMV appeared more evident when using biotinylated versus radiolabeled Vip3Aa protein. This suggests that binding properties are better preserved when biotinylating versus radiolabeling Vip3Aa. Similarly, radiolabeling of Cry1F affected specific binding (57), while specific binding was still detected using biotinylated Cry1F (58). Therefore, disparities in Vip3Aa binding parameters and binding site alteration between studies may be due to a combination of binding conditions and the insect species tested.

Resistance to Vip3Aa in *C. virescens* was found to associate with reduced membrane-bound ALP expression and corresponding enzymatic activity compared to a susceptible strain (35). However, this ALP protein did not function as a receptor for Vip3Aa in a heterologous system. Lack of differences in ALP and APN activity in CEW-Vip-RR compared to CEW-SS supports that these BBMV proteins are not involved in the resistance mechanism.

In conclusion, the results of this study provide evidence that a Vip3Aa receptor is altered in a resistant strain of *H. zea*. This observation explains lack of cross-resistance to Cry proteins and predicts cross-resistance to multiple Vip3A proteins sharing binding sites in CEW-Vip-RR. Based on this observation and widespread *H. zea* resistance to Cry1 and Cry2 proteins (9–13), new PIPs not sharing binding sites with Vip3Aa would be ideal for mitigating the negative consequences and spread of Vip3Aa resistance. Caution should be taken when deciding which Bt traits are used in regions where *H. zea* is likely to develop Vip3A resistance. Current research efforts are focused on transcriptomic and proteomic analyses in identifying crucial receptors for Vip3Aa toxicity altered in CEW-Vip-RR.

## MATERIALS AND METHODS

### Insect strains

The Vip3Aa susceptible (CEW-SS) and resistant (CEW-Vip-RR) *H. zea* strains were established as previously described (17, 18, 59). The CEW-SS strain was obtained from Benzon Research Inc. (Carlisle, PA) in 2018 and has documented susceptibility to Cry1Ac, Cry2Ab, and Vip3Aa (59, 60). The CEW-Vip-RR strain was established in 2019 from an F_2_ screen of gravid female moths collected by light trapping in Snook, TX (18). Diet overlay bioassays most recently showed that CEW-Vip-RR had a Vip3Aa resistance ratio of >45,100-fold relative to a susceptible strain (17). Resistance to Vip3Aa in CEW-Vip-RR was confirmed to follow an autosomal monogenic recessive pattern of inheritance (17). Susceptible and resistant strains were made near-isogenic through four rounds of repeated crossing, backcrossing, and reselection with a discriminatory dose of Vip3Aa protein (3.16 µg/cm^2^).

### Protein production and purification

The full-length Vip3Aa39 protein used in this study was produced in a recombinant *Escherichia coli* strain previously described (18). An overnight preculture was inoculated in LB media containing 100 mg/ml of ampicillin and incubated (37°C, 180 rpm) until reaching an OD_600_ of 0.6-0.8. The culture was then induced with 1 M isopropyl-β-d-thiogalactopyranoside (IPTG) and incubated overnight. Bacterial cells were collected via centrifugation (15,000 x *g*, 8 min, 4°C) and the pellets were resuspended in lysis buffer (50 mM Na_2_CO_3_, 0.1 M NaCl, 3 mg/ml lysozyme, 10 µg/ml DNAse I). The sample was then sonicated on ice for seven cycles (5 sec on/5 sec off, amplitude 65%) and left overnight at 4°C with mild shaking. The soluble Vip3Aa protoxin was collected in the supernatant after centrifugation (17,500 x *g*, 20 min, 4° C) and purified via anion-exchange chromatography with an AKTA pure chromatography system (GE Healthcare). The sample was passed through an anion-exchange column (HiTrap Q HP) equilibrated with buffer A (50 mM Na_2_HCO_3_, 50 mM NaHCO_3_; pH 9.8), and then eluted in fractions using a step gradient of buffer B (buffer A containing 1 M NaCl). SDS-PAGE was used to confirm the presence of Vip3Aa in fractions corresponding to the elution peak. Purified samples containing Vip3Aa were pooled and the concentration of Vip3Aa was determined using the Qubit protein assay kit (Invitrogen) prior to storage at -80°C.

For experiments requiring Vip3Aa toxin, protoxin was processed by incubation with 0.2 mg/ml of trypsin (Sigma, ≥10,000 BAEE units/mg) for three hours at 37°C prior to purification by anion-exchange chromatography as above. The Cry1Ac protein was produced in the HD73 strain of *B. thuringiensis* (obtained from the *Bacillus* Genetic Stock Center, Columbus, OH), and purified as described elsewhere (40).

### Larval gut pH measurements

Fourth-instar *H. zea* larvae were starved overnight and subsequently divided into four treatment groups of nine larvae each (each larva representing a biological replicate). The first group was fed with a 5 µl droplet of milli-Q water containing 50 mg/ml sucrose and a small amount of bromophenol blue to help visualize droplet ingestion. The other groups were fed with droplets containing 5 µg of Vip3Aa protoxin, 10 µg of Vip3Aa activated toxin, or 5 µg of Cry1Ac protoxin. Doses were selected because they were capable of killing susceptible larvae within one day of feeding. After 2 hours of ingestion, larvae were immobilized with stainless steel pins and their gut exposed by cutting a longitudinal incision along the dorsal midline of the cuticle. A microelectrode (Thermo Scientific, CN: 9863BN) was used to measure the pH in five sections of the gut (foregut, anterior midgut, central midgut, posterior midgut, and hindgut), as previously described (40).

### Protease extraction and preparation

The midguts of actively feeding fourth-instar *H. zea* larvae were dissected to extract soluble gut proteases. The food bolus from ten guts (representing one biological replicate) were pooled together in a microcentrifuge tube, flash frozen in liquid nitrogen and stored at -80°C until used. Soluble gut proteases were extracted by thawing the gut contents on ice before adding 500 µl of milli-Q water, followed by vortexing for two minutes. The tubes were then centrifuged at 27,000 x *g* for 20 minutes at 4°C and the supernatants filtered with 0.22 µm filters to obtain the soluble protease sample. Protein concentrations in soluble midgut protease extracts were estimated using a Qubit protein assay kit and adjusted to 1 mg/ml before storing at -80°C.

### Insecticidal protein processing

Processing of Vip3Aa protoxin (80 µg) by 7 µg of soluble midgut protease extracts was tested in a final volume of 140 µl of buffer (50 mM Na_2_HCO_3_, 50 mM NaHCO_3_; pH 9.8) at 37°C with mild shaking. Aliquots (5 µl) were taken after 10, 30, 60, 120, 240, and 360 minutes and also after 21 hours, and reactions were stopped by adding one volume of 2X sample buffer (61). The samples were heat denatured for 10 minutes before storing at -20°C. Cry1Ac protoxin processing was also tested using the same methods with the exception that 5 µg of midgut fluid extracts were added to protoxin, and aliquots were taken at 5, 10, 15, 20, 30, 45, and 60 minutes. Samples were resolved in AnykD Criterion Precast SDS-PAGE gels (Bio-Rad) and stained for total protein using ProtoBlue Safe (National Diagnostics). Protoxin processing was quantified by densitometry with ImageJ software (62) by measuring the intensity of the remaining protoxin band in each lane relative to the amount of input protoxin.

### BBMV preparation and enzyme activity assays

Midguts were dissected from fourth-instar *H. zea* larvae and immediately frozen in liquid nitrogen prior to storage at -80°C. Brush border membrane vesicles (BBMV) were prepared from frozen midguts using differential magnesium centrifugation (63), and final BBMV pellets were suspended in ice-cold phosphate-buffered saline (PBS) buffer (137 mM NaCl, 2.7 mM KCl, 1.5 mM KH_2_PO_4_, 1 mM Na_2_HPO_4_; pH 7.4). Final BBMV protein concentrations were determined using a Qubit protein assay kit (Invitrogen) and then used for aminopeptidase N (APN) and alkaline phosphatase (ALP) enrichment tests as previously described (64). Activity was assessed in two separate BBMV preparations tested in triplicate for each preparation by comparing initial midgut homogenates and final BBMV samples using leucine *p*-nitroanilide and *p*-nitrophenyl phosphate as substrates for APN and ALP activity, respectively. A BioTek Synergy H1 plate reader (Agilent) with Gen5 software (Agilent) was used to record enzymatic activities and calculate Max V.

### Biotinylation and radiolabeling of Vip3Aa toxin

Vip3Aa toxin was biotinylated with a 1:30 (protein:biotin) molar ratio of EZ-Link Sulfo-NHS-LC-Biotin (Life Technologies) for 30 minutes on ice. Excess biotin was removed from samples via dialysis against PBS buffer for two hours at room temperature followed by dialysis overnight at 4°C.

Purified Vip3Aa (25 µg) and Cry1Ac (1 µg) toxins were radiolabeled with 0.5 mCi of NaI^125^ (Perkin Elmer) using chloramine T, as described before (65). Specific activities were 0.27 mCi/pmol for Vip3Aa and 1.34 mCi/pmol for Cry1Ac.

### Vip3Aa and Cry1Ac binding assays

Homologous competition binding assays were performed by incubating 30 μg of BBMV proteins with 45 nM of biotinylated Vip3Aa in 0.1 ml (final volume) of binding buffer (PBS, 0.1% Tween-20, 0.1% bovine serum albumin) for one hour at 25°C. Non-specific binding was determined from reactions that included a 100-fold molar excess of unlabeled Vip3Aa as a competitor. Reactions were stopped by centrifugation (27,000 x *g*, 10 min, RT) and the resulting pellets were washed with 0.5 ml of ice-cold binding buffer. The centrifugation and washing process was repeated and final pellets were solubilized in 15 μl of 1X electrophoresis sample buffer (60). Samples were heat denatured for five minutes and then resolved in SDS-10% PAGE gels. Resolved proteins were electro-transferred (20 V) to 0.45 μm nitrocellulose membranes (Thermo Scientific) in transfer buffer (192 mM glycine, 25 mM Tris, 0.1% SDS [w/v], 20% methanol; pH 8.3) at 4°C overnight. Membranes were blocked for 45 minutes in PBST buffer (137 mM NaCl, 2.7 mM KCl, 1.5 mM KH_2_PO_4_, 1 mM Na_2_HPO_4_, 0.1% Tween-20, pH 7.4) with 3% BSA for 1 hour, and then probed with a 1:30,000 dilution of streptavidin-HRP in the blocking buffer. Biotinylated toxin was visualized using a SuperSignal West Pico PLUS Chemiluminescent Substrate kit (Thermo Scientific), according to the manufacturer’s instructions, in an Amersham Imager 600 (GE Healthcare). Two biological replications (different BBMV preparations) each with two technical replications were tested. The intensity of the Vip3Aa toxin band was estimated using densitometry and specific binding was calculated by subtracting non-specific binding from total binding (in the absence of competitor) values. Each blot was standardized by dividing the value for specific binding by the value corresponding to the band intensity of 10 ng of biotinylated Vip3Aa toxin.

Saturation and competition binding assay reactions were run with 20 µg of BBMV proteins and ^125^I-Vip3Aa as a ligand in 0.1 ml (final volume) reactions in binding buffer. In saturation assays, total binding was determined from reactions containing up to 15 nM of ^125^I- Vip3Aa, and non-specific binding was determined from reactions that included a 100-fold molar excess of unlabeled Vip3Aa. Competitive binding assays were performed in reactions containing a fixed concentration of ^125^I-Vip3Aa (12 nM) or ^125^I-Cry1Ac (1.1 nM) and increasing concentrations of unlabeled homologous competitor. All saturation and competition assay reactions were performed as described above for homologous competition assays. The amount of radioactivity in the final pellets was measured with a WIZARD^2^ gamma counter (Perkin Elmer). All binding assays were replicated using two different BBMV preparations each with two to three technical replications. Results from competition assays were analyzed using KELL software (Biosoft, Cambridge, United Kingdom) to estimate equilibrium disassociation constants and binding site concentrations.

### Statistical analyses

Larval gut pH measurements were rank-transformed and analyzed using a two-way repeated measure analysis of variance (ANOVA) using SAS/STAT v. 9.4 (SAS Institute Inc., Cary, NC). Strain and treatment were treated as between-subject effects, whereas gut regions were treated as a within-subject effect. A Tukey-Kramer post-hoc test with α of 0.05 was used for mean separation. After rank transformation, processing of Vip3Aa and Cry1Ac was analyzed using a generalized linear mixed model including processing time and strain as fixed effects.

Random effects included biological replications. A Tukey post-hoc test with an α of 0.5 was used for mean separation. An independent t-test was used to evaluate binding of biotinylated Vip3Aa to BBMV from CEW-SS and CEW-Vip-RR using JMP Pro v. 16 (SAS Institute Inc., Cary, NC). Specifically bound ^125^ I-Vip3Aa in saturation assays was analyzed using a generalized linear model in SAS/STAT v. 9.4 with fixed effects including strain and input ^125^ I-Vip3Aa. A Tukey-Kramer post-hoc test with an α of 0.5 was used for mean separation. Independent t-tests run in JMP Pro v. 16 were used to compare the enzymatic activities of midgut homogenates and BBMVs from CEW-SS and CEW-Vip-RR larvae.

## ACKNOWLEDGEMENTS

The authors thank Xiaocun Sun for assistance in statistical analysis. This research was funded by grant 2021-67013-33567 from the Agriculture and Food Research Initiative program of the USDA National Institute of Food and Agriculture.

